# A CARF-HAD phosphatase effector provides immunity during the type III-A CRISPR-Cas response

**DOI:** 10.1101/2025.04.30.651354

**Authors:** Gianna Stella, Linzhi Ye, Sean F. Brady, Luciano Marraffini

**Affiliations:** Laboratory of Bacteriology, The Rockefeller University, New York, NY 10065, USA; Tri-Institutional PhD Program in Chemical Biology, Weill Cornell Medical College, Rockefeller University and Memorial Sloan Kettering Cancer Center, New York, NY 10065, USA; Laboratory of Genetically Encoded Small Molecules, The Rockefeller University, New York, NY 10065, US; Howard Hughes Medical Institute, The Rockefeller University, New York, NY 10065, USA

## Abstract

CRISPR-Cas systems provide adaptive immunity against phage infection in prokaryotes using an RNA-guided complex that recognizes complementary foreign nucleic acids. Different types of CRISPR-Cas systems have been identified that differ in their mechanism of defense. Upon infection, Type III CRISPR-Cas systems employ the Cas10 complex to find phage transcripts and synthesize cyclic oligo-adenylate (cOA) messengers. These ligands bind and activate CARF immune effectors that cause cell toxicity to prevent the completion of the viral lytic cycle. Here we investigated two proteins containing an N-terminal haloacid dehalogenase (HAD) phosphatase domain followed by four predicted transmembrane helices and a C-terminal CARF domain, which we named Chp. We show that, in vivo, Chp localizes to the bacterial membrane and that its activation induces a growth arrest, leads to a depletion of ATP and IMP and prevents phage propagation during the type III CRISPR-Cas response. In vitro, the CARF domain of Chp binds cyclic tetra-adenylates and the HAD phosphatase domain dephosphorylates dATP, ATP and IMP. Our findings extend the range of molecular mechanisms employed by CARF effectors to defend prokaryotes against phage infection.

## INTRODUCTION

Prokaryotes have evolved multiple defense strategies to fight invading genetic elements such as plasmids and bacteriophages (1). <u>C</u>lustered <u>R</u>egularly Interspaced <u>S</u>hort <u>P</u>alindromic <u>R</u>epeats (CRISPR) loci, and <u>C</u>RISPR-<u>as</u>sociated (*cas*) genes provide RNA-guided, adaptive immunity to bacteria and archaea (2). Depending on the *cas* gene content, CRISPR-Cas systems can be classified into different types (3). Type III systems employ the multi-protein Cas10 complex, which harbors an RNA guide known as the CRISPR RNA (crRNA) (4). Upon infection, annealing of the crRNA with a complementary RNA sequence present in a transcript produced by the invader (5) leads to the stimulation of three distinct activities of the Cas10 complex: (i) non-specific ssDNA degradation by the Cas10 HD nuclease domain (6), (ii) conversion of ATP into cyclic-oligoadenylate (cOA) second messengers by the Cas10 Palm cyclase domain (7, 8), and (iii) cleavage of the complementary target RNA (5). Cleavage of ssDNA intermediates which are generated during phage and plasmid DNA transcription and replication are thought to directly destroy the invader’s genome (9). During phage infection, this activity mediates effective immunity when the target transcript is expressed early during the lytic cycle (9, 10), presumably due to the presence of a low number of phage genomes. Synthesis of cOAs, primarily cyclic tetra- and hexa-adenylates (cA_4_ and cA_6_), activates type III immune effectors that contain a <u>C</u>RISPR <u>a</u>ssociated <u>R</u>ossman <u>F</u>old (CARF) domain to bind these second messengers (7, 8). These effectors usually harbor a second domain whose enzymatic activity is stimulated by the binding of the cOA ligand and disrupts the host metabolism to interfere with the completion of the phage lytic cycle and limit viral spread (4). The activation of CARF effectors is fundamental to provide immunity to the bacterial population when the target transcript is expressed late in the lytic cycle and the ssDNase activity of the Cas10 complex is not sufficient to eliminate the phage DNA from the host (9, 11–13), and results in a growth arrest of the infected cells (11–14). Finally, cleavage of the target transcript ends both of these activities (5, 6), temporarily stopping the type III CRISPR-Cas response until another intact target transcript is recognized by the crRNA of the Cas10 complex.

Type III CRISPR loci encode many different CARF effectors, with a remarkable functional and structural diversity (4). To date, the activities of CARF effectors include ssDNA and ssRNA degradation (11, 15–18), membrane depolarization (12), ATP deamination (13, 19) and NAD^+^ cleavage (20). Here, we investigated the role of a CARF effector containing a haloacid dehalogenase (HAD) phosphatase domain previously identified by a bioinformatic search of novel genes associated with CRISPR loci (20). HAD phosphatases belong to the HAD hydrolase superfamily, one of the largest known enzyme classes, present in all three kingdoms of life (21, 22). HAD superfamily members contain a highly conserved α/β core domain with four conserved motifs (I-IV), and in many cases also a cap domain that acts as an active site lid to sterically constrain the size and shape of its substrates (23–25). HAD phosphatases catalyze the cleavage of P-OP bonds (26), but due to the relatively low sequence conservation across different enzymes, predicting substrate specificity is challenging. For example, one study that probed *in vitro* the substrate specificity of 23 soluble HAD phosphatases encoded in the *E. coli* genome against 80 phosphorylated metabolites revealed varying degrees of substrate promiscuity (27). The most common substrates were phosphorylated carbohydrates, pyridoxal 5’-phosphate (PLP), flavin mononucleotide (FMN), and nucleotides. Among HAD phosphatases that hydrolyzed nucleotides, YrfG preferentially hydrolyzed purines (GMP and IMP), YjjG hydrolyzed pyrimidines (UMP, dUMP, dTMP) and YieH hydrolyzed both purines and pyrimidines, although its primary substrate is glucose 6-phosphate (27). *In vivo*, YjjG has been determined to function as a house-keeping nucleotidase that recognizes noncanonical nucleobase derivatives and prevents their misincorporation into DNA (28). Another characterized HAD phosphatase, NagD, showed activity against UMP and GMP (29). Outside of prokaryotic organisms, HAD phosphatases hydrolyze nucleotides in both plants (30) and mammals (31, 32). The cellular function of many HAD hydrolases remains unknown and it was proposed that their broad substrate specify may serve as a reservoir for evolutionary novelty (27). We studied the role of the <u>C</u>ARF-<u>H</u>AD <u>p</u>hosphatase effectors, which we named Chp1 and Chp2, in the type III-A CRISPR-Cas immune response of staphylococci. We found that they localized to the staphylococcal membrane and, when activated by cA_4_, induce a growth arrest that protects bacteria against phage infection, most likely through the hydrolysis of ATP and dATP.

## RESULTS

### Chp1 reduces ATP and IMP levels to mediate cell toxicity during the type III-A CRISPR-Cas response

A previous bioinformatic search for novel CRISPR-associated genes (20) led to the identification of two genes encoding an N-terminal HAD phosphatase domain followed by four predicted transmembrane helices and a C-terminal CARF domain. We named these genes CRISPR-associated HAD phosphatase 1 and 2 (*chp1* and *chp2*; Figs 1A-B). A structural prediction of Chp1 made using AlphaFold3 (33) suggested the formation of a dimer through the interaction between the CARF domains of two Chp1 monomers (Fig. 1C). In support of this prediction, all CARF effectors which structures have been determined to date have a dimeric architecture to generate a symmetric binding site for cA_4_ or cA_6_ (4). Additional contacts between the α-helices of two different subunits seem to reinforce the dimeric structure. In contrast, HAD phosphatase domains are not predicted to interact within the dimer. Chp2 shares 84% sequence similarity with Chp1 (Fig. S1A) and is predicted to have an equivalent structure (Fig. 1D).

**Figure 1.**
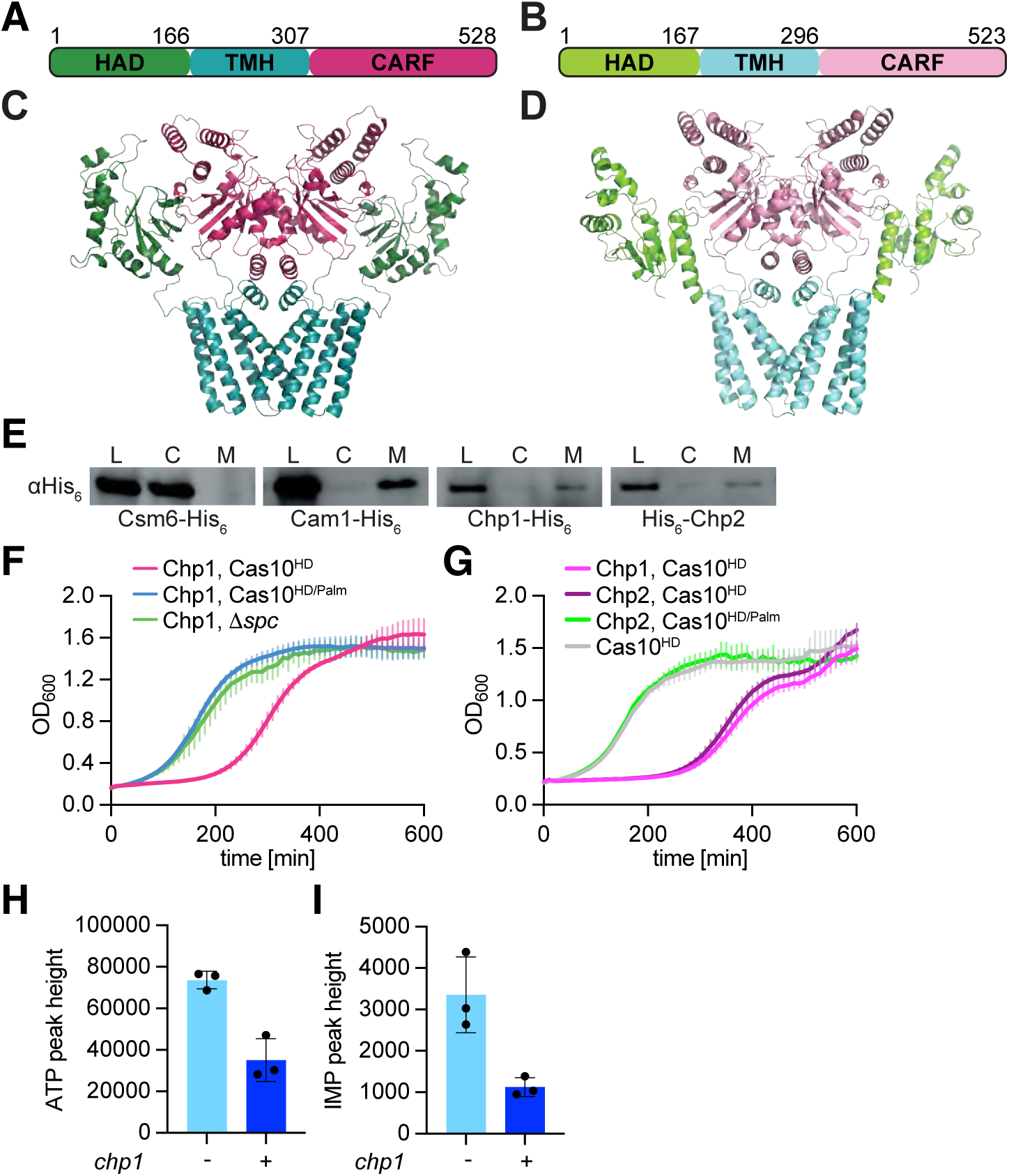
Activation of Chp1 leads to cell toxicity and reduced ATP and IMP levels in vivo. **(A)** Domain architecture of *Bacteriodales* Chp1 which was identified within a type III-D CRISPR-*cas* locus. **(B)** Domain architecture of *Prevotella sp*. Chp2 which was identified within a type III-A CRISPR-*cas* locus. **(C)** Alphafold3 structure of *Bacteriodales* Chp1. The protein contains a N-terminal HAD phosphatase domain (green) followed by transmembrane helices (blue) and a C-terminal CARF domain (pink). Residue numbers are labeled. **(D)** Alphafold3 structure of *Prevotella* Chp2. The protein contains a N-terminal HAD phosphatase domain (light green) followed by transmembrane helices (light blue) and a C-terminal CARF domain (light pink). Residue numbers are labeled. **(E)** Western blot of cellular fractions. Staphylococci harboring plasmids encoding hexahistidine-tagged Csm6, Cam1, Chp1, and Chp2 were outgrown and lysed. Total lysate fraction (L) were collected and then subjected to ultracentrifugation to obtain cytosolic (C) and membrane (M) fractions. Samples were blotted with a primary anti-His_6_ antibody. **(F-G)** Growth of staphylococci carrying pTarget and pCRISPR variants, measured as OD_600_ after the addition of aTc. Data are mean of three biological triplicates ± SEM (H) Quantification of ATP and (I) IMP levels from bacterial lysates. Extracts from staphylococci harboring pTarget and pCRISPR(DChp1) or pCRISPR(Chp1) were collected after 15 minutes of incubation with aTc and analyzed via LC-MS. Mean of three biological replicates ± SEM is reported. The *p* values shown were obtained with a two-sided t test with Welch’s correction.

The *chp1* gene is located within a type III-D CRISPR-*cas* locus from an unidentified *Bacteriodales* bacterium (Fig. S1B). Similarly, *chp2* exists within a type III-A locus present in *Prevotella sp.* (Fig. S1B). To investigate the role of these CARF effectors in the type III CRISPR-Cas response, we cloned each of them individually into pCRISPR, a plasmid harboring the *Staphylococcus epidermidis* RP62a type III-A CRISPR-*cas* locus (34), which is genetically related to the type III systems that naturally harbor *chp1* and *chp2* (Fig. S1B). In this plasmid we replaced the open reading frame encoding for the staphylococcal CARF effector Csm6 with either of the *chp* genes, generating pCRISPR(Chp) (Fig. S1C). Using this construct to express Chp1 and Chp2 in staphylococci, we tested their subcellular localization through the introduction of a C- and N-terminal hexa-histidyl tag, respectively. Expression of C-terminally tagged Csm6 and Cam1 served as cytoplasmic and membrane-bound controls, respectively (12). We obtained staphylococcal lysates expressing these different CARF effectors and performed an anti-hexa-histidine western blot (Fig. 1E and S1D) and found that both Chp1 and Chp2 primarily associate with the cell membrane, most likely via the predicted transmembrane helices.

We previously investigated the function of CARF effectors through the evaluation of cell toxicity and growth arrest induced upon activation of the Cas10 complex by a target RNA (11–14, 20). We therefore generated a *S. aureus* RN4220 strain (35) harboring both pCRISPR(Chp1) and pTarget (14). pTarget encodes a target RNA under a tetracycline-inducible promoter. The Cas10 complex is expressed from pCRISPR(Chp1) and was programmed with a crRNA complementary to target RNA. To prevent pTarget degradation by the nuclease activity of the Cas10 complex (14), we introduced mutations in the HD domain (H14A, D15A) of the Cas10 subunit, generating pCRISPR(Chp1, Cas10^HD^). To test for Chp1 toxicity, we treated cultures with anhydro-tetracycline (aTc) to induce target RNA expression and production of cOAs by the Cas10 complex and followed bacterial growth by measuring optical density at 600 nm (OD_600_). We found that, compared to control cultures that do not produce cOAs due to the absence of either a crRNA that can recognize the target (Chp1, Δ*spc*) or cOA synthesis (mutations in the cyclase active site of Cas10; D586A,D587A; Cas10^Palm^), the growth of staphylococci carrying Chp1 was delayed (Fig. 1F). This growth arrest depended on the presence of the CARF effector, since it was not detected in cells harboring a plasmid that lacks a CARF effector [pCRISPR(Cas10^HD^)] (Fig. 1G). A test of *chp2* in similar genetic backgrounds showed that this CARF-HAD phosphatase variant also generates a growth arrest upon production of cOAs during the staphylococcal type III-A CRISPR-Cas response (Fig. 1G).

Given that some of the most common substrates of HAD phosphatases are NTPs (27), and that a common anti-phage defense strategy is the alteration of the nucleotide pool of the host to prevent viral propagation (36–39), we decided to investigate the effect of Chp1 activation on the abundance of different nucleotides within staphylococci, using untargeted liquid chromatography-mass spectrometry (LC/MS) analysis. We collected cells from cultures either expressing [pCRISPR(Cas10^HD^, Chp1)] or lacking [pCRISPR(Cas10^HD^, Δ*chp1*)] Chp1, in which cOA production was induced by the addition of aTc for 15 minutes, and quantified 27 different nucleotides in each sample. We observed that Chp1 activation resulted in a significant decrease in ATP (Fig. 1H) and IMP levels (Fig. 1I) but not for other nucleotides (Fig. S2). Together this data demonstrates that Chp1 activation during the type III-A CRISPR-Cas response leads to a reduction of ATP and IMP levels and cause a growth arrest in staphylococci. We hypothesized that this is a consequence of (i) the binding of cOA second messengers to the CARF domain of the Chp effectors that (ii) stimulate the nucleotide phosphatase activity of the HAD domain. We therefore performed biochemical experiments to test these hypotheses.

### The CARF domains of Chp1 and Chp2 bind cA_4_

To determine whether Chp1 and Chp2 bind any of the cOAs produced by the cyclase activity of the Cas10 RNA-guided complex, cA_3_, cA_4_ and cA_6_ (7, 8), we decided to purify both proteins using heterologous expression in *Escherichia coli*. However, most likely due to the presence of the transmembrane helices, we found the proteins to be insoluble. We therefore expressed and purified the N-terminal hexa-histidyl tagged CARF domains of Chp1 and Chp2 (Fig. 2A). We then performed thermal shift assays, mixing the purified CARF domains (5 μM) with 100 μM of either cA_3_, cA_4_, cA_6_ and a fluorescent dye that binds to exposed hydrophobic residues to monitor protein thermal stability, and followed the change in fluorescence during heating of the mixture. We calculated the first derivative of relative fluorescence units [-d(RFU)/dT] to determine the protein melting temperature values (T_m_) in the presence of each cOA (Fig. 2B). We found that the T_m_ of Chp1-CARF was 41°C either in the absence of ligand or in the presence of cA_3_, and 42°C in the presence of cA_6_, indicating these cOAs do not stabilize the protein. In contrast, when Chp1-CARF was incubated with cA_4_ the T_m_ shifted to 75°C, an observation that demonstrates that tetra-adenylate is the ligand for Chp1. Similar results were obtained with purified Chp2-CARF (Fig. 2A), with a T_m_ of 51°C either in the absence of ligand or in the presence of cA_3_, 52°C in the presence of cA_6_, but 70°C when incubated with cA_4_ (Fig. 2C). Finally, we calculated the binding affinity of Chp1-CARF for its ligand, by incubating 5 μM of the protein with varying concentrations of cA_4_ and fluorescent dye and monitoring the change in fluorescence upon heating. We then normalized the -d(RFU)/dT measurements obtained at 41°C (the T_m_ of apo Chp1-CARF) for the different ligand concentrations (Fig. 2D). As a consequence of the shift of the T_m_ to 75 °C upon ligand binding, the normalized -d(RFU)/dT values decreased as the cA_4_ concentration increased. We performed a nonlinear regression to fit a binding curve to our data, which was used to determine the equilibrium dissociation constant (K_d_) of 273 nM for the binding of cA_4_ to Chp1-CARF. These data demonstrates that the CARF domains of both Chp1 and Chp2 bind cyclic tetra-adenylates.

**Figure 2.**
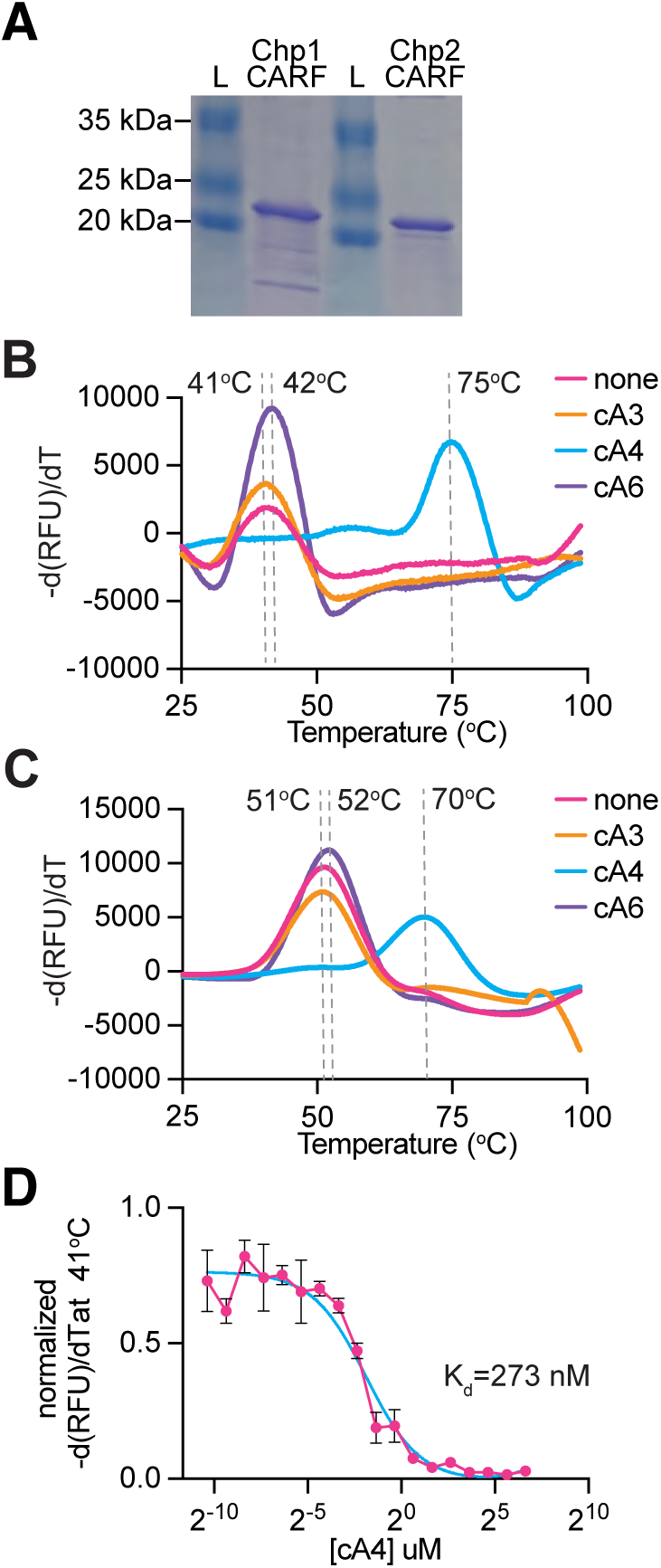
The Chp1 and Chp2 CARF domains bind cA_4_. **(A)** SDS-PAGE stained with Coomassie showing purified Chp1 CARF domain and purified Chp2 CARF domain. **(B)** Thermal shift assay of purified Chp1 CARF domain (5 uM) incubated with cAn ligand (100 uM) or no ligand. First derivative of Relative Fluorescence Units plotted as a function of temperature. Mean of three technical triplicates is reported. **(C)** Same as **(B)** but with purified Chp2 CARF domain. Mean of two technical duplicates is reported. **(D)** Thermal shift assay of purified Chp1 CARF domain (5 uM) incubated with varying concentrations of cA_4_. The first derivative of Relative Fluorescence Units at 41°C was normalized and then plotted as a function of cA_4_ concentration. A nonlinear regression for one site binding (saturation) was performed to calculate K_d_. The fit line is in cyan. Mean of three technical triplicates, ±SEM, is reported.

### Chp1 dephosphorylates ATP and IMP

Given the reduction in ATP and IMP levels that we detected *in vivo* upon activation of Chp1 and Chp2 (Fig. 1H-I), and that nucleotides are common substrates of HAD phosphatases (27), we decided to test for the enzymatic activity of Chp1 and Chp2 on a diverse set of nucleotides. To do this, we purified an N-terminal hexa-histidyl tagged version of the HAD phosphatase domain of Chp2 (Fig. 3A). Unfortunately, in spite of using different affinity tags and purification protocols, we were not able to obtain a protein preparation of the HAD phosphatase domain of Chp1 suitable for biochemical studies. We incubated Chp2-HAD (50 μM) with ATP, IMP, UTP, CTP, GTP, TTP, dCTP, dGTP or dATP (1 mM), Mn^+2^ (1 mM), in the presence or absence of EDTA, and separated the reaction products using HPLC. We were unable to detect changes for UTP, CTP, GTP, TTP, dCTP nor dGTP (Fig. S4A-E). In contrast, we observed conversion of dATP to dADP (32%), ATP to ADP (12%) and IMP to inosine (22%) (Figs. 3B-D). EDTA prevented this conversion, a result that indicates that Mn^+2^ is required for the reaction, similar to other HAD phosphatases (27, 40). This experiment demonstrates that Chp2 has phosphatase activity *in vitro*. In addition, degradation of ATP and IMP correlates with the *in vivo* LC/MS results obtained after activation of Chp1 *in vivo*, for which we observed a decrease in the cellular levels of these two nucleotides (Fig. 1H-I).

**Figure 3.**
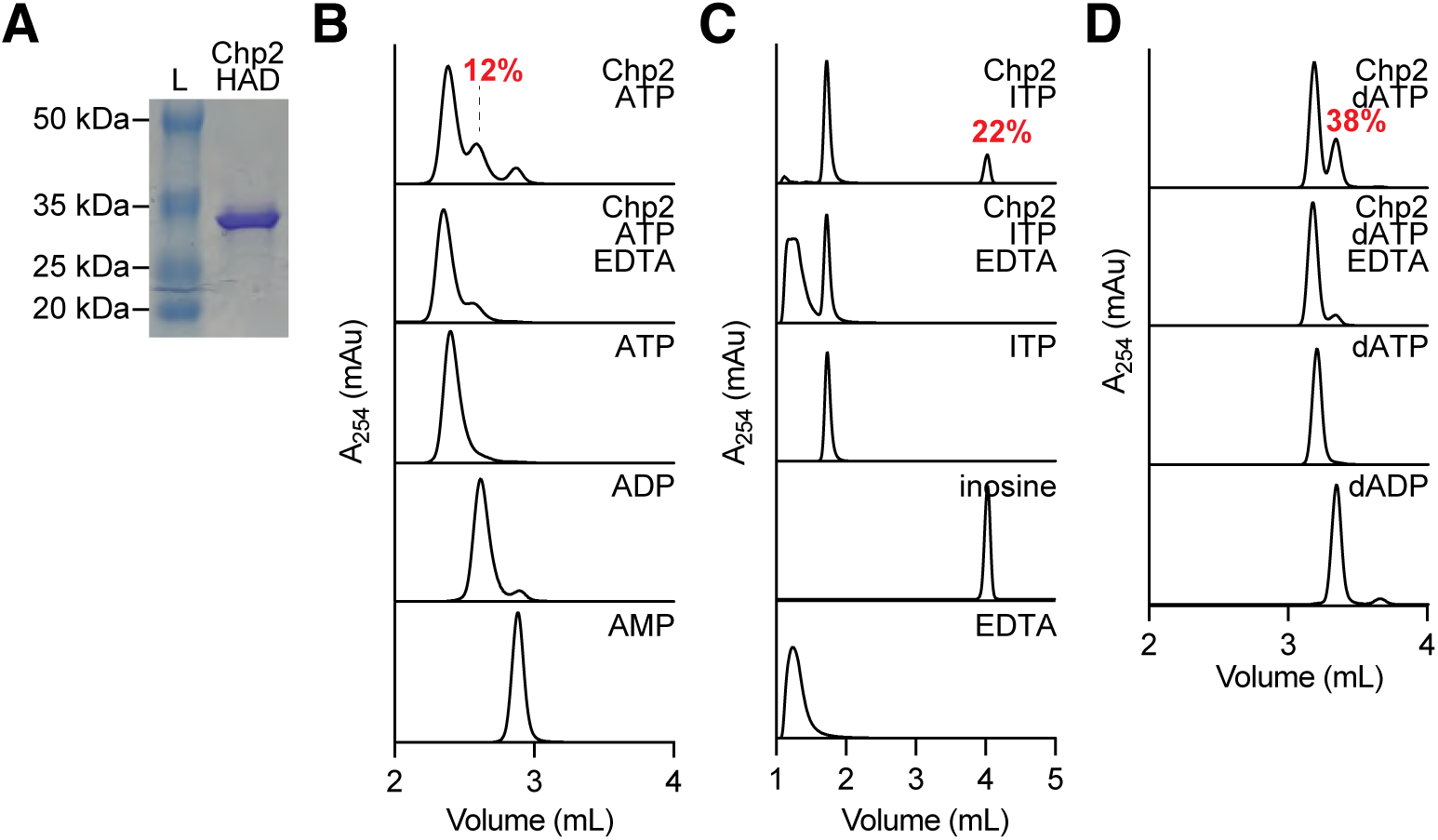
Chp2 dephosphorylates ATP, IMP, and dATP. **(A)** SDS-PAGE stained with Coomassie showing purified Chp2 HAD domain. **(B)** HPLC analysis of Chp2 (50 μM) reaction products in the presence ATP (1 mM). Chromatograms of ATP, ADP, and AMP are shown as standards. Reactions were performed in duplicate. Percentage of substrate was calculated using quantified using peak integrations of reaction with or without EDTA. Average percentage from two duplicates is shown. **(C)** Same as with **(B)** but in the presence of IMP (1 mM). Chromatograms of IMP, inosine, and EDTA are shown as standards. **(D)** Same as with **(B)** but in the presence dATP (1 mM). Chromatograms of dATP and dADP are shown as standards.

### Conserved residues found in HAD phosphatases are not essential for Chp1 toxicity

A previous bioinformatic analysis of HAD hydrolases indicated that although they share little overall sequence similarity (15-30%), they contain four conserved motifs, I-IV {refs Koonin}. Motif I contains a signature DxD sequence, where the aspartic acids coordinate the metal cofactor required for hydrolysis. The first aspartate acts as a nucleophile and forms an aspartyl-intermediate during catalysis (41, 42). In the case of HAD phosphatases, the second aspartate acts as a general acid-base. Motifs II and III contain highly conserved threonine or serine and lysine, respectively, and may contribute to the stability of the reaction intermediates during hydrolysis (43). Motif IV contains acidic residues with the signatures D<u>D</u>, G<u>D</u>xxxD, or G<u>D</u>xxxxD, where the underlined aspartate is highly conserved and also contributes to the coordination of the metal cofactor (42, 44). We aligned Chp1 and Chp2 to ten HAD hydrolase sequences that were previously characterized bioinformatically using MUSCLE (45), and found that they contain motifs I, II and IV, and a minimal motif III (Fig. 4A). Moreover, Chp1 contains all five highly conserved residues: D8, D10, S75, K118, and D129. A structural prediction of the HAD phosphatase domain of Chp1 in the presence of ATP made using AlphaFold3 (33) suggested that all these residues reside in a metal ion pocket (Fig. 4B). Chp2 also contains the same conserved residues (Fig. 4A) forming a similarly predicted pocket (Fig. S4A). To test the importance of these residues for Chp1 activity, we introduced alanine substitutions to generate the following mutants: D8A/D10A, D8A/K118A and D8A/K118A/D129A. We then tested Chp1’s ability to induce a growth arrest in staphylococci during the activation of type III-A CRISPR-Cas immunity (Figs. S4B, 4C). Surprisingly, we found that none of the substitutions prevented the growth delay caused by wild-type Chp1. We therefore made deletions that encompass the entire HAD phosphatase domain for both CARF effectors, generating Chp1(Δ2-146) and Chp2(Δ2-138), which in both cases eliminated Chp toxicity (Fig. S4C). While we were not able to identify the active site residues, this result suggests that the HAD phosphatase domain is responsible for Chp1 and Chp2 activity.

**Figure 4.**
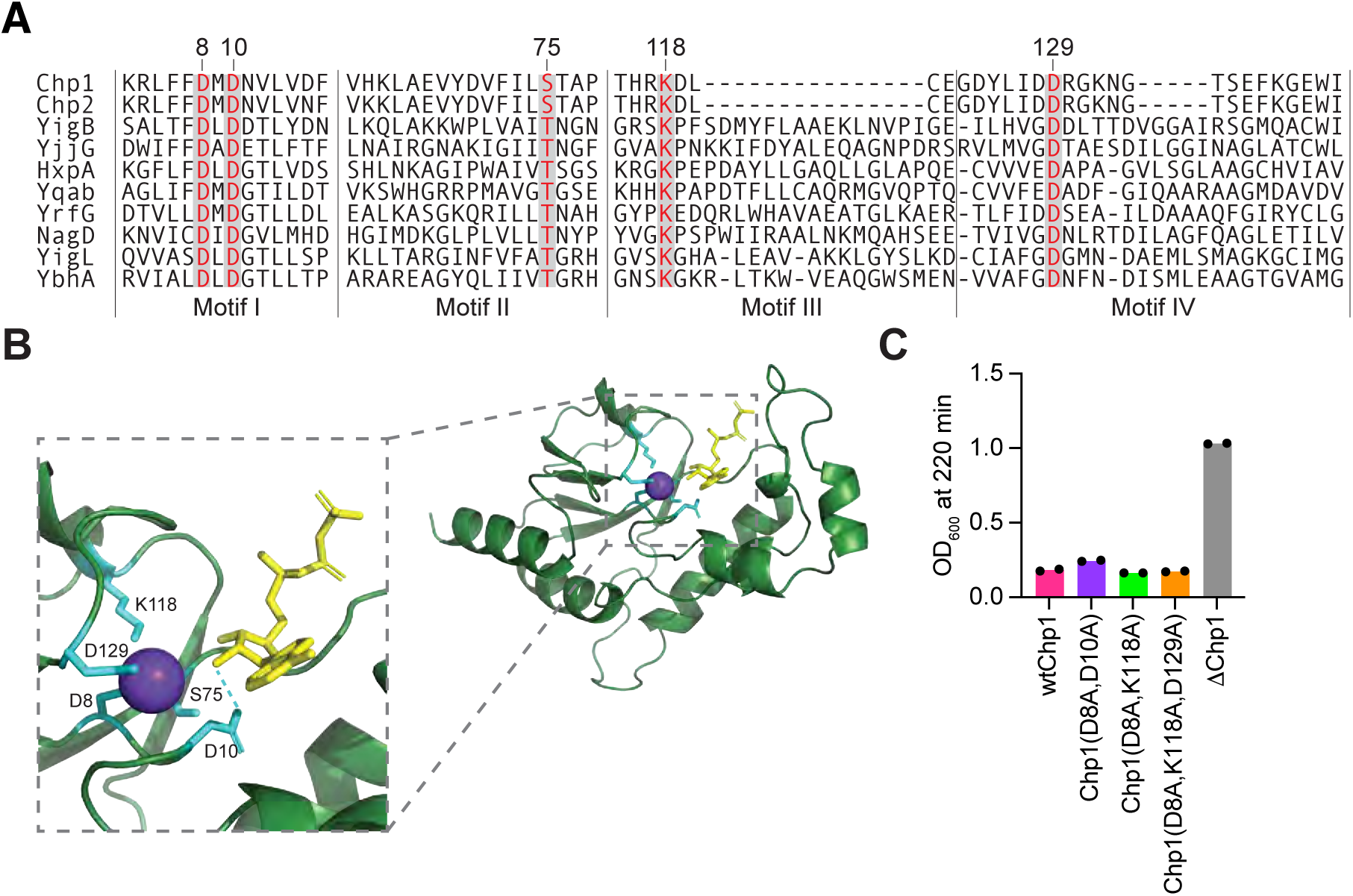
Chp1 contains conserved residues found in HAD phosphatases. **(A)** Protein alignment of known motifs from E. coli HAD phosphatases with Chp1 and Chp2. Highly conserved residues are highlighted. Numbering for Chp1 residues is shown. **(B)** Alphafold3 structure of HAD phosphatase domain from Chp1 with conserved residues noted (cyan). Modeled with ATP (yellow) and Mn^2+^ (purple). **(C)** Growth of staphylococci harboring pTarget and pCRISPR(Chp1) with mutations in highly conserved HAD residues measured as OD_600_ value at 220 min after addition of aTc. Data are mean of two biological duplicates ± SEM.

### Chp1 provides anti-phage defense

For the majority of CARF effectors studied to date, the toxicity they mediate is essential to provide anti-phage immunity when the target transcript is expressed late during the viral lytic cycle and Cas10’s nuclease activity is not sufficient to restrict infection (9–13, 20). We tested the function of Chp1 in the type III-A CRISPR-Cas response after infection with the staphylococcal phage ϕNM1γ6, at multiplicity of infection (MOI) 1 and 5. We programmed the Cas10 complex with crRNA guides that recognize either the early- or late-transcripts of the *gp14* or *gp43* genes, respectively (the crRNAs originated from *spc14* and *spc43*, respectively) (Fig. 5A). As reported before, immunity mediated by *spc14* did not require the Chp1 effector but depended on the nuclease activity of Cas10, both at MOI 1 (Fig. 5B) and 5 (Fig. 5C). In the absence of Cas10 nuclease activity, however, Chp1 alone was able to support immunity at MOI 1 (Fig. 5B) but not at 5 (Fig. 5C), consistent with an abortive infection mode of defense. Interestingly, the activation of Chp1 early during infection generated a noticeable growth delay at MOI 1, which provides further evidence that Chp1 induces growth arrest during phage infection. In contrast, Chp1 was sufficient to provide defense mediated by *spc43* at MOI 1, with staphylococci proliferating with a marked delay (Fig. 5D). At MOI 5, Chp1 alone did not enable the survival of the bacterial culture, and required the nuclease activity of Cas10 (Fig. 5E), a similar result to that obtained with other CARF effectors heterologously expressed in staphylococci such as Cam1 (12) and Cad1 (13). These results were corroborated by enumerating plaque forming units (PFU) three hours after infection with ϕNM1γ6 at MOI 1 (Fig. 5F). We found that, when type III-A CRISPR immunity is triggered late in the phage lytic cycle via *spc43*, Chp1, but not Cas10, alone prevented the increase in PFU values observed in the absence of CRISPR immunity (Δ*spc*). In contrast, the presence of both Chp1 and the nuclease activity of Cas10 caused a marked decrease of viral propagation. To investigate these results at the cellular level, we performed fluorescence microscopy, taking advantage of GFP expression by ϕNM1γ6^GFP^ to visualize infected cells (9). Consistent with the growth arrest mediated by Chp1 activation that we observed in the presence of both pTarget (Fig. 1F) and phage (Fig. 5D), infection of cultures carrying a type III-A CRISPR-Cas system programmed *spc43* and expressing Chp1 and Cas10^HD^ led to a stop in the division of the infected cells (marked by green fluorescence) that was followed by the division of non-infected ones (non-fluorescent) (Fig. 5G). In contrast, in the absence of Chp1, the green infected cells lysed. Based on these data, we conclude that the growth arrest generated by Chp1 affects the ability of ϕNM1γ6 to complete its infectious cycle and the release of viral particles from infected hosts, allowing non-infected cells to proliferate. Finally, we tested the ability of Chp1 carrying mutations in conserved HAD phosphatase residues, (D8A/D10A, D8A/K118A and D8A/K118A/D129A) to provide anti-phage defense. In line with the results obtained for Chp1 toxicity assays, we found that none of the mutations affected immunity against ϕNM1γ6 infection, measured as a reduction in PFUs (Fig. 5H).

**Figure 5.**
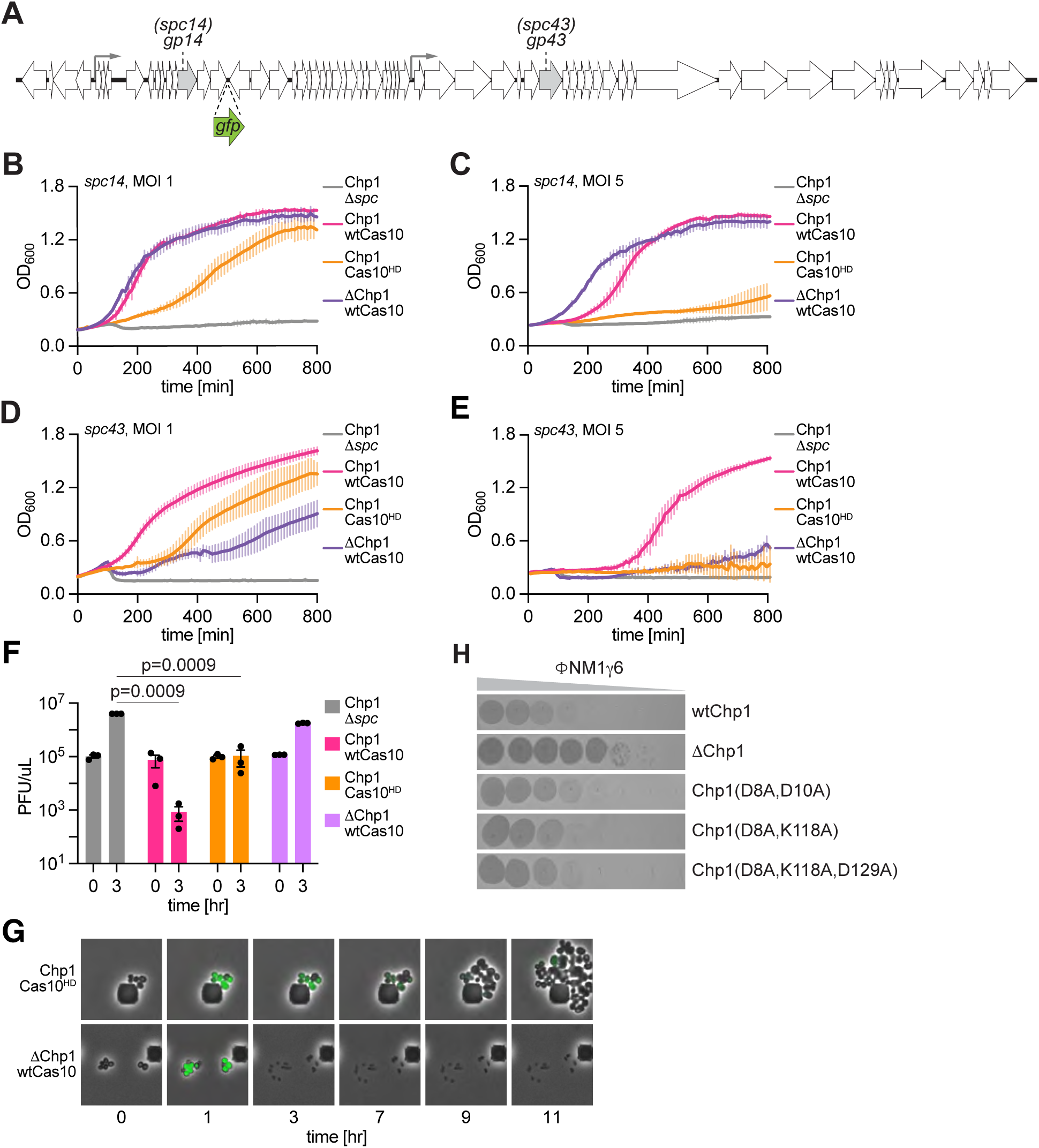
Chp1 is required for anti-phage defense. **(A)** Schematic of the genome of staphylococcal phage ΦNM1γ6 showing the location of the transcripts targeted by different spaces of the type III-A CRISPR-Cas system. Gray arrows indicate promoters. GFP is inserted at indicated location for fluorescent microscopy experiments. **(B)** Growth of staphylococci carrying different pCRISPR constructs with a spacer to target the *gp14* transcript of ΦNM1γ6. Measured OD_600_ with ΦNM1γ6 at an MOI of 1. Mean of three biological triplicates, ±SEM, is reported. **(C)** same as **(B)** at an MOI of 5. **(D)** same as **(B)** but targeting the *gp43* transcript. **(E)** same as **(D)** but at an MOI of 5. **(F)** Enumeration of plaque-forming units within staphylococcal cultures harboring different pCRISPR constructs with a spacer to target the *gp43* transcript at the indicated times after infection with ΦNM1γ6 at an MOI of 1. The mean of three biological replicates, ±SEM, is reported. The *p* values shown were obtained with a two-sided t test with Welch’s correction. **(G)** Time course microscopy of *S. aureus* encoding different pCRISPR constructs after infection with ΦNM1γ6-GFP. Images are representative of two biological replicates. **(H)** Plaques from phage ΦNM1γ6 on *S. aureus* lawns with pCRISPR(Chp1, Cas10^HD^) where constructs contain either mutant or WT Chp1. Negative controls include Cas10^HD/dPalm^ or Δ*spc* CRISPR.

## DISCUSSION

Here we investigated the role of CARF effectors harboring HAD phosphatase domains in the staphylococcal type III-A CRISPR-Cas immune response. We found that two highly similar homologs, Chp1 and Chp2, localize to the bacterial membrane, and, upon activation of the Cas10 complex by a target transcript, bind cA_4_ cyclic nucleotides and cause HAD phosphatase-dependent cytotoxicity to prevent viral propagation. The impossibility of purifying full-length Chp1 or Chp2, most likely due to the presence of the transmembrane helices, prevented us from demonstrating a direct stimulation of the phosphatase activity after cA_4_ binding. We have two reasons to believe this is the case. First, although we were not able to test cyclic oligoadenylate binding by the full-length proteins, previous studies have shown an excellent correlation of the ligand specificity and binding between purified CARF domains and full-length effectors (13). Second, we determined that cell toxicity is triggered by cA_4_ synthesis *in vivo* and depends on the presence of the HAD phosphatase domain of Chp1 and Chp2. Based on these observations, we propose that ligand binding by the CARF domain results in a conformational change that stimulates the HAD phosphatase activity of Chp1 and Chp2.

How these CARF effectors cause toxicity and provide immunity is less clear. Several bacterial anti-phage defense systems deplete nucleotides to halt the lytic cycle and prevent viral propagation. For example, cytidine-deaminases deplete dCTP by converting it to dUTP (37, 39), effectors associated with <u>c</u>yclic-oligonucleotide-<u>b</u>ased <u>a</u>nti-phage <u>s</u>ignaling <u>s</u>ystems (CBASS) possess ATP nucleosidase activity that cleaves this fundamental nucleotide into adenine and ribose-5’-triphosphate (38), and GajB, the effector of the Gabija system, has phosphatase activity that hydrolyzes and depletes ATP, GTP, dATP, and dGTP (36). Given these precedents, that some of the most common substrates of HAD phosphatases are NTPs (27), and that our *in vitro* and *in vivo* data demonstrated NTP dephosphorylation and nucleotide depletion, respectively, we hypothesized that Chp1 and Chp2 provide defense through the hydrolysis of phosphate groups from nucleotides, leading to a decrease in the cellular levels of these important metabolites. More specifically, we observed dephosphorylation of ATP, dATP and IMP *in vitro* and detected depletion of ATP and IMP, but not dATP, *in vivo*. dATP is generated directly and indirectly from ATP molecules by ATP reduction by ribonucleotide reductases (46) or phosphorylation of dAMP and dADP using ATP as the donor of phosphate groups by nucleoside kinases (47), respectively. Therefore, we believe that it is possible that Chp1 and Chp2 also dephosphorylate dATP *in vivo*, but that the reduction of dATP is compensated by consuming ATP molecules; a scenario consistent with our results. Importantly, we did not explore the effects of Chp1 and Chp2 activation on other common HAD phosphatase substrates such as pyridoxal 5’-phosphate (PLP), and flavin mononucleotide (FMN) (27). In principle, degradation of these metabolites could cause the reduction in cell growth and phage propagation we detected. Therefore, it remains a possibility that phosphorylated substrates other than nucleotides are additional targets of Chp1 and Chp2 during the type III-A CRISPR-Cas response. This is a necessary consideration given that our in vitro assays were performed with only the HAD phosphatase domain of Chp2, outside of the membrane localization context and cA_4_-mediated activation.

Whether by depleting nucleotides or other phosphorylated metabolites, Chp1 and Chp2 activation during the type III-A response generates a marked growth arrest that is important to provide immunity against phage infection. Like other CARF effectors, this mode of defense is most important when the target transcript is expressed late in the viral lytic cycle and in the absence of Cas10 ssDNase activity. This is especially important because approximately only 40% of type III CRISPR-Cas systems contain variants that either lack an HD domain (48) or do not display detectable ssDNase activity (49). When compared with other CARF effectors, Chp1 and Chp2 are unique in that they cause ATP depletion and therefore it is intriguing to think that their activity may decrease cyclic oligoadenylate synthesis, which requires ATP {ref}, and negatively impact the immune response they mediate. CARF effectors display a great diversity of mechanisms to halt viral propagation, which include ssDNA and ssRNA degradation (11, 15–18), membrane depolarization (12), ATP deamination (13, 19) and NAD^+^ cleavage (20). By incorporating nucleotide depletion to this broad list of activities, Chp1 and Chp2 expand the range and flexibility of the type III CRISPR-Cas immune response.

## Acknowledgements.

We would like to thank Adrian Morales Amador for help with the LC/MS protocols. LAM is supported by funds from NIH R01GM149834. LAM is an investigator of the Howard Hughes Medical Institute.

## Author contributions

LAM and GS conceived the study. GS performed all experiments except LC/MS, which was carried out by LY under the supervision of SB. The manuscript was written by LAM and GS.

## Competing interests

LAM is a cofounder and Scientific Advisory Board member of Intellia Therapeutics, a cofounder of Eligo Biosciences and a Scientific Advisory Board member of Ancilia Biosciences.

## Methods

### Sequence alignments

Alignments and calculations of sequence identity and similarity were determined either using MUSCLE (45) or VectorBuilder. Alignments for Figure 4 were manually corrected to ensure conserved residues were aligned.

### Bacterial growth

*Staphylococcus aureus* strain RN4220 (50) was grown at 37°C in brain heart infusion (BHI) medium supplemented with 10 μg ml^−1^ chloramphenicol to maintain pCRISPR and 10 μg ml^−1^ erythromycin to maintain pTarget. 5 μM CaCl_2_ was supplemented in phage experiments unless indicated otherwise.

### Plasmid cloning

The plasmids and oligonucleotides used in this study are detailed in Table S1. The amino acid sequences of Chp1 and Chp2 were sourced from NCBI GenBank (reference sequences CADBDX000000000.1 from *Bacteriodales* bacterium and GCA_017627465.1 from *Prevotella* sp., respectively) and synthesized by Azenta.

### Chp1 toxicity assay

To measure the effect of Chp1 activity on *S. aureus* growth over time, colonies of *S. aureus* containing pTarget and the specified pCRISPR were launched in liquid culture overnight in triplicate. The next day, cells were diluted 1:100 and then normalized for optical density at a wavelength of 600 nm (OD_600_) after an 1 hour and 15-minute outgrowth. Next, pTarget transcription was induced using 12.5 ng mL^-1^ aTc. OD_600_ readings were obtained every 10 minutes with a microplate reader.

### Quantification of phage plaques

To quantify plaque-forming units (PFU) over time from cultures infected with phage, *S. aureus* cultures containing various pCRISPRs were launched overnight, diluted 1:100 and outgrown for about 1 hour. Then, cells were infected with phage ΦNM1g6 at an MOI of 1. Aliquots were taken from infected cells at the beginning of infection, 1 hour, and 3 hours post-infection. These aliquots were spun down to remove cells, subject to 10-fold serial dilutions, and spotted onto RN4220.

### Time-course fluorescence microscopy of phage-infected cultures

To visualize the dynamics of phage infection and immunity provided by Chp1, colonies of *S. aureus* containing various pCRISPRs with spacers programmed to target ϕNM1γ6-GFP were observed under phase contrast and in a GFP fluorescence channel as previously described (12, 13).

### Cell fractionation and western blotting

To determine localization of Chp1 and Chp2, cultures were fractionated and subjected to western blotting. RN4220 cultures containing hexahistidine-tagged Csm6, Cam1, Chp1, or Chp2 were launched overnight. All overnight culture were diluted to OD_600_ of 0.05 and outgrown for 2 hours. Cells were spun down at 3,900 rpm, decanted, and resuspended in lysis buffer (20 mM HEPES pH 7.1, 150 mM NaCl, 10% glycerol).

Resuspended cultures were incubated with 2 mg/mL lysostaphin and a protease inhibitor cocktail, cOmplete Tablets EDTA-free EASYpack (Roche) at 37°C for 15 minutes. Lysates were then sonicated and spun down 3,900 rpm. An aliquot of the supernatant was collected as a whole cell lysate sample. The remaining supernatant was then subjected to ultracentrifugation at 100,000 g for 1 hour using a TLA-120.2 rotor. An aliquot of the supernatants was collected as a cytosolic sample and the remainder was discarded. The membrane pellets were resuspended in resuspension buffer and homogenized using a Teflon Douce homogenizer. The homogenized pellets were then subjected to another round of ultracentrifugation at 100,000 g for 1 hour. The supernatants were removed, and the pellets were resuspended once more in resuspension buffer using a Teflon Douce homogenizer. An aliquot was saved as a membrane sample. These samples were then run on a 4-20% Mini-PROTEAN TGX Precast Protein Gels (Bio-Rad). Transferred proteins were then probed with THE^TM^ His Tag Antibody, mAb, Mouse (1:5,000 dilution, GenScript A00186). Goat anti-Mouse IgG (H+L) Secondary Antibody, HRP (1:10,000 dilution, Thermo Fisher 31430) was used to prepare the blot for imaging.

### Protein expression and purification

An N-terminal 6xHis tagged Chp1 CARF domain (307–528), an N-terminal 6xHis tagged Chp2 HAD domain (1–138), and an N-terminal 6xHis Chp2 CARF domain (295–523) were independently cloned into a pET28 vector under an IPTG-inducible promoter and transformed into Escherichia coli strain BL21 (DE3). Bacteria were grown at 37 °C to an OD_600_ of 0.5 – 0.7 and induced with 0.5 mM isopropyl β-d-1-thiogalactopyranoside (IPTG) at 18 °C overnight. Bacterial cells were spun down, resuspended in lysis buffer (20 mM HEPES pH 7.3, 400 mM NaCl, 30 mM imidazole, 10% glycerol) supplemented with protease inhibitor cocktail, cOmplete Tablets EDTA-free EASYpack (Roche), and sonicated. Cell lysates were centrifuged at 11,000 rpm for 10 minutes. Supernatants were filtered using a 0.22 mm PES filter and applied to a gravity column with pre-equilibrated nickel resin. The column was washed with wash buffer 1 (20 mM HEPES pH 7.3, 1 M NaCl, 30 mM imidazole, 10% glycerol), wash buffer 2 (same as lysis buffer), and then protein was eluted with elution buffer (20 mM HEPES pH 7.3, 400 mM NaCl, 300 mM imidazole, 10% glycerol). Fractions containing the protein of interest were combined and dialyzed into 20 mM HEPES pH 7.3, 150 mM NaCl, 10% glycerol.

### Thermal shift assays

To determine the melting temperature (T_m_) of the CARF domains of Chp1 and Chp2, a thermal shift assay was performed using kit 4462263 from ThermoFisher. 5 uM of purified Chp1 (307–528) or Chp2 (295–523) was co-incubated with either 100 uM of cA_3_, cA_4_, cA_6_, or no ligand. Fluorescence was measured using the Melt Curve program on the QuantStudio 3 Real-Time PCR System (Applied Biosystems) as the mixture was heated at a rate of 0.1 °C/s from 25°C to 98°C. The first derivative of the Relative Fluorescence Units (RFU) was plotted over temperature to determine T_m_ for each condition. To determine the dissociation constant (K_d_), 5 uM of purified Chp1 (307–528) was co-incubated with 2-fold dilutions of cA_4_ (from 100 uM to 0.76 nM) and heated as described above. The first derivatives of the Relative Fluorescence Units (RFU) at 41°C, the apo T_m_, were normalized against each other from 0 to 1 using the equation: (RFU – min RFU)/(max RFU – min RFU). These values were then plotted against cA_4_ concentration and fit using a nonlinear regression for one site binding to determine the K_d_ value.

### Analysis of common nucleotides and nucleosides using HPLC-coupled high-resolution mass spectrometry

Colonies containing pTarget and pCRISPR plasmids were launched overnight. The following day, cultures were diluted 1:100, grown to OD_600_ of 0.3, and induced with 125 ng/mL aTc for 15 minutes before pelleting and collection. Cells were resuspended in 4 mM EDTA dissolved in 80% ethanol and vortexed vigorously. Next, cells were heated to 85°C with shaking for 3 minutes and allowed to cool to room temperature. Cell lysates were spun down at 13,000 rpm in a tabletop centrifuge and supernatants were then further processed. About 600 uL of supernatant was dried under blowing air at room temperature for 2 hours. The remaining extracts were reconstituted to 100 uL with 10 mM ammonium bicarbonate solution (pH 7) and run through 10 kDa filters at 14,000 x g, 4°C for 30 minutes. The filtrates and chemical standards of common nucleotides and nucleotides were analyzed on Sciex X500 QTOF mass spectrometer with electrospray ionization in the negative mode. Samples were delivered by ExionLC AC system equipped with ACQUITY UPLC HSS T3 column (2.1 x 100 mm, 1.8 μm) that was held at 40°C. Solvent A was water with 10 mM ammonium acetate (pH 6.9) and solvent B was acetonitrile. Flow rate was 0.5 ml/min. For each run, 1 uL of sample was injected and separated with the following gradient: 0% B for 1.5 min, to 30.0% B in 3.5 min, to 95% B in 1.0 min, and 95% B for 1.0 min. Column eluate between 0.5 min and 5 min was directed to MS using the following method: curtain gas of 30 psi, ion source gas 1 of 50 psi, ion source gas 2 of 50 psi, temperature of 400°C, soray voltage of 550. For TOFMS, m/z between 100 and 1000 was scanned with a declustering potential of 80 V, collision energy of 10 V, collision gas of 7 psi, and accumulation time of 0.1 second. For TOFMSMS, MRM (Multiple Reaction Monitoring) was applied: declustering potential of 80 V, collision energy of 35 V, accumulation time of 0.03 seconds. All data were collected and analyzed on the software Sciex OS. The levels of common nucleotides and nucleosides were semi-quantified using the peak intensity of each molecule’s characteristic MRM transition.

### In vitro dephosphorylation reactions

Dephosphorylation reactions were performed *in vitro* by incubating substrates at 37°C for 90 minutes. Reaction substrates were incubated at a final concentration of 1 mM with 50 uM Chp1 (1–168) or Chp2 (1–138) HAD domains in reaction buffer (20 mM HEPES pH 7.1, 150 mM NaCl, 1 mM TCEP, 1 mM MnCl_2_ and 10% glycerol). After incubation at 37°C, reactions were quenched by heating mixtures to 65°C for 5 minutes. Reaction mixtures were diluted with nuclease-free water to 100 uL before being filtered with Amicon® Ultra Centrifugal Filter, 10 kDa MWCO filters to remove proteins before analysis. 10 uL of filtered reaction products were then injected onto an Agilent Bonus-RP, 4.6 x 150 mm, 3.5 um Rapid Res. C18 column held at 40°C at a flow rate of 1.2 mL per min. For ATP, ADP, AMP, IMP, Inosine, dATP, and dADP, the following mobile phase buffer was used in the elution program: 60 mM K_2_HPO_4_, 40 mM KH_2_PO_4_, pH 7.0. For GTP, GDP, CTP, CDP, UTP, and, UDP, 10 mM TBAB was supplemented into the mobile phase buffer. The elution program was as follows: 0 min 100 % buffer, 0 % ACN; 2 min 95 % buffer, 5 % ACN; 4 min 80 % buffer, 20 % ACN; 5.3 min 75 % buffer, 25 % ACN and 6 min 100 % buffer, 0 % ACN. Chromatograms were collected by monitoring absorbance at 254 nm. To determine substrate conversion, integrated peaks from reaction mixtures without EDTA were compared to integrated peaks from reaction mixtures with EDTA.

---

**Figure S1.**
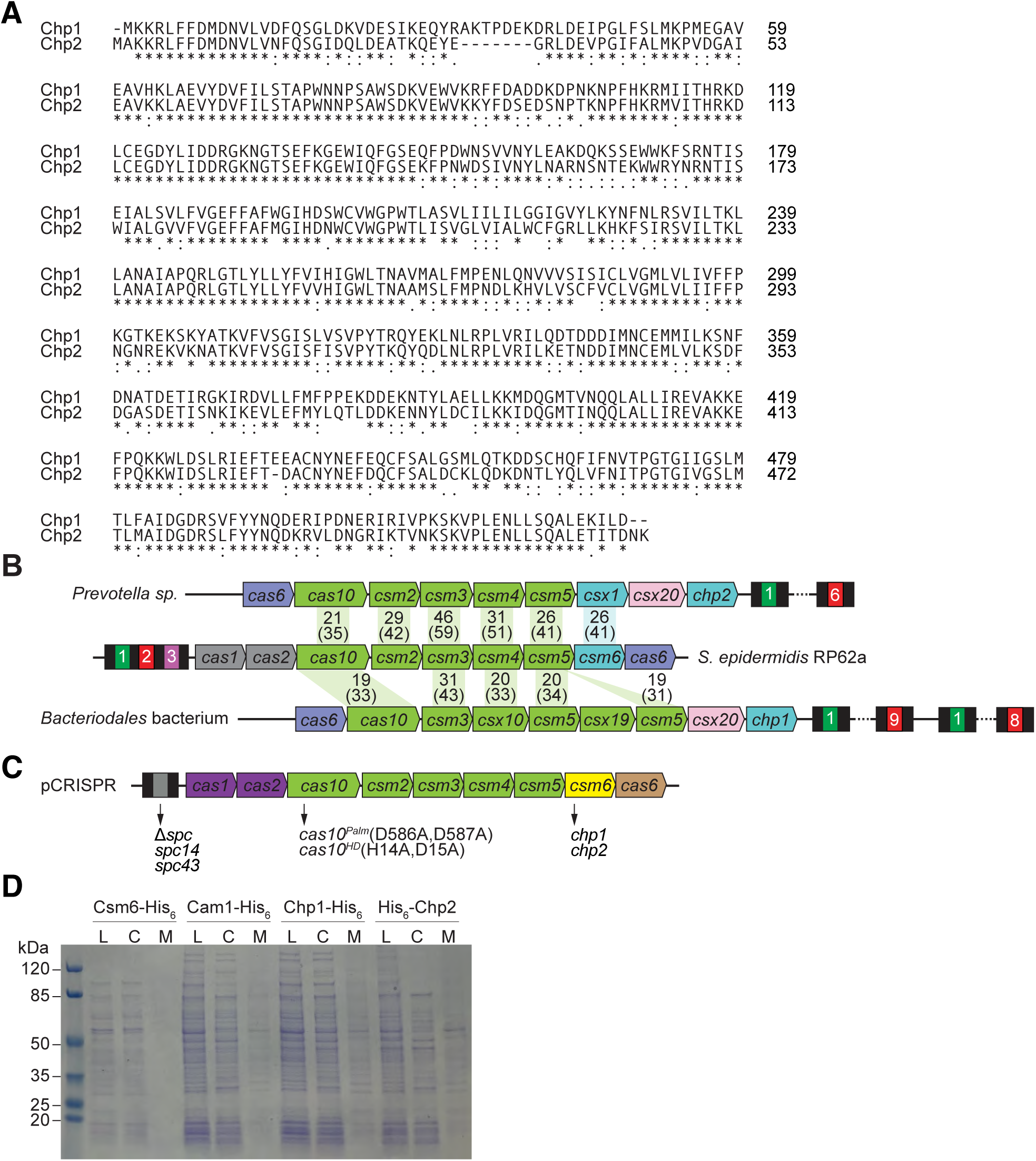
**(A)** Clustal Omega sequence alignment of full-length Chp1 and full-length Chp2 (*) indicates identical residues; (:) indicates strongly similar residues; (.) indicates weakly similar residues. **(B)** Comparison of the III-A systems of *Prevotella sp*. and *Staphylococcus epidermidis* RP62a and the type III-D system of *Bacteriodales* bacterium. Black boxes indicate CRISPR repeats; colored, numbered boxes indicate CRISPR spacers. Numbers indicate the percent homology of amino acid sequences. Numbers in parentheses indicate the percent homology at the DNA sequence level. **(C)** Genetic modifications of the *S. epidermidis* RP62a type III-A CRISPR locus cloned into various pCRISPR plasmids. Amino acid substitutions, domain deletions, and insertion of different spacer sequences are indicated. **(D)** SDS-PAGE with Coomassie staining showing lysate, cytosolic, and membrane fractions used for western blotting experiment in Figure 1E.

**Figure S2.**
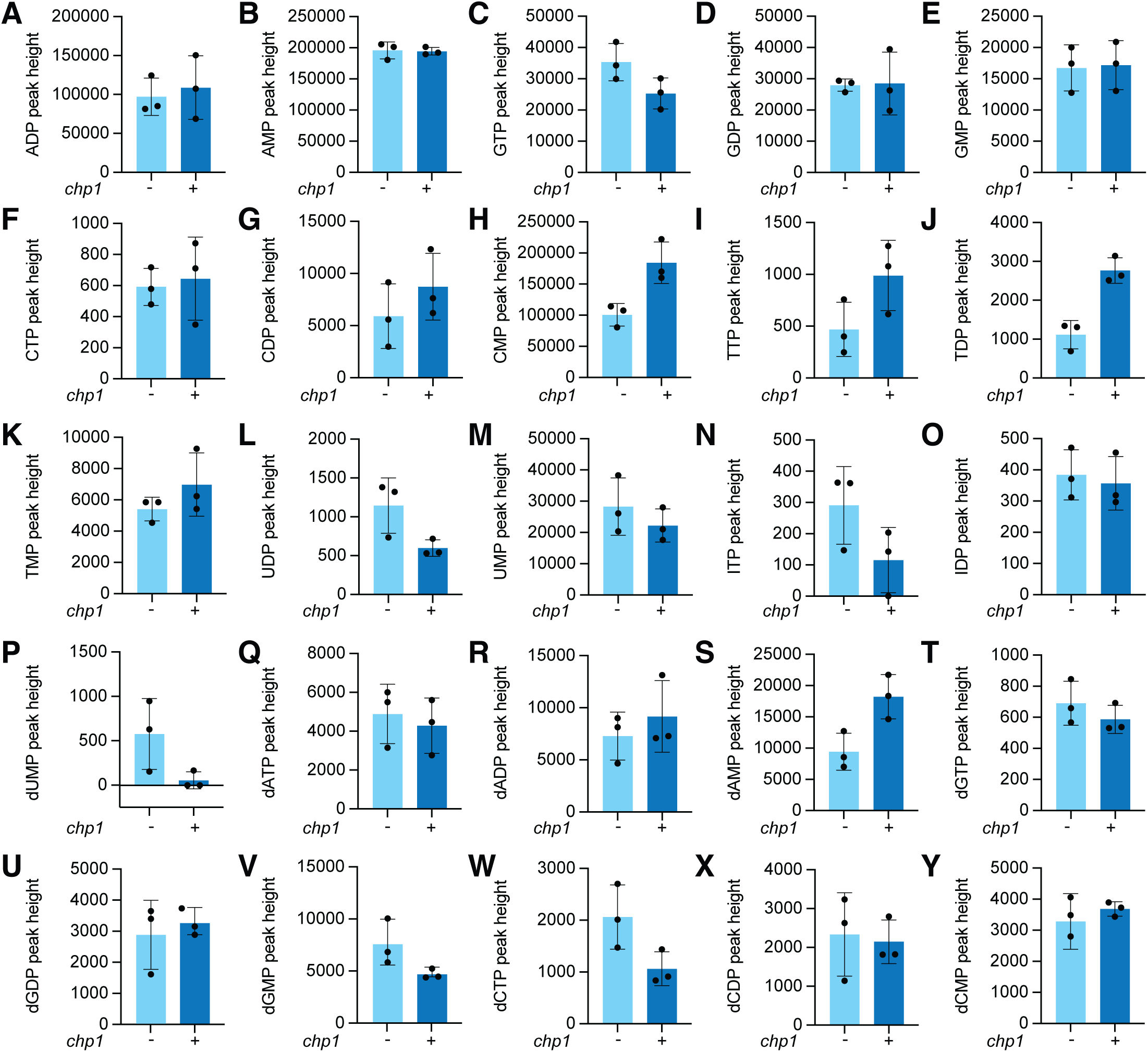
**(A-Y)** Quantification of various nucleotide levels from bacterial lysates. Extracts from staphylococci harboring pTarget and pCRISPR(ΔChp1) or pCRISPR(Chp1) were collected after 15 minutes of incubation with aTc and analyzed via LC-MS. Mean of three biological replicates ± SEM is reported.

**Figure S3.**
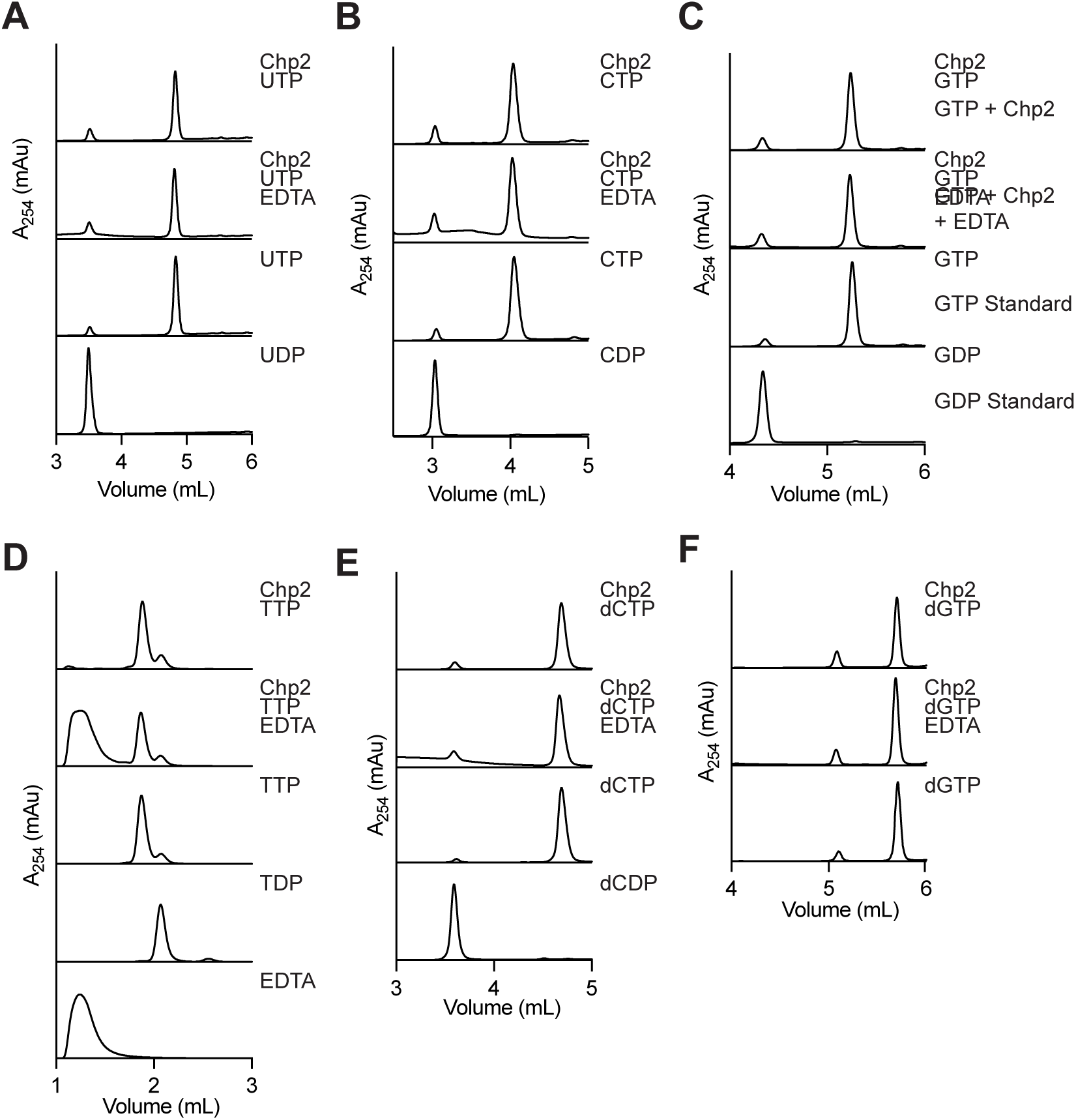
**(A)** HPLC analysis of Chp2 (50 μM) reaction products in the presence UTP (1 mM). Chromatograms of UTP and UDP are shown as standards. Reactions were performed in duplicate. **(B)** Same as **(A)** but in the presence CTP (1 mM). Chromatograms of CTP and CDP are shown as standards. Reactions were performed in duplicate. **(C)** Same as (A) but in the presence GTP (1 mM). Chromatograms of GTP and GDP are shown as standards. Reactions were performed in duplicate. **(D)** Same as **(A)** but in the presence TTP (1 mM). Chromatograms of TTP and TDP are shown as standards. Reactions were performed in duplicate. **(E)** Same as **(A)** but in the presence of dCTP (1 mM). Chromatograms of dCTP and dCDP are shown as standards. Reactions were performed in duplicate. **(F)** Same as **(A)** but in the presence of dGTP (1 mM). Chromatograms of dGTP and dGDP are shown as standards. Reactions were performed in duplicate.

**Figure S4.**
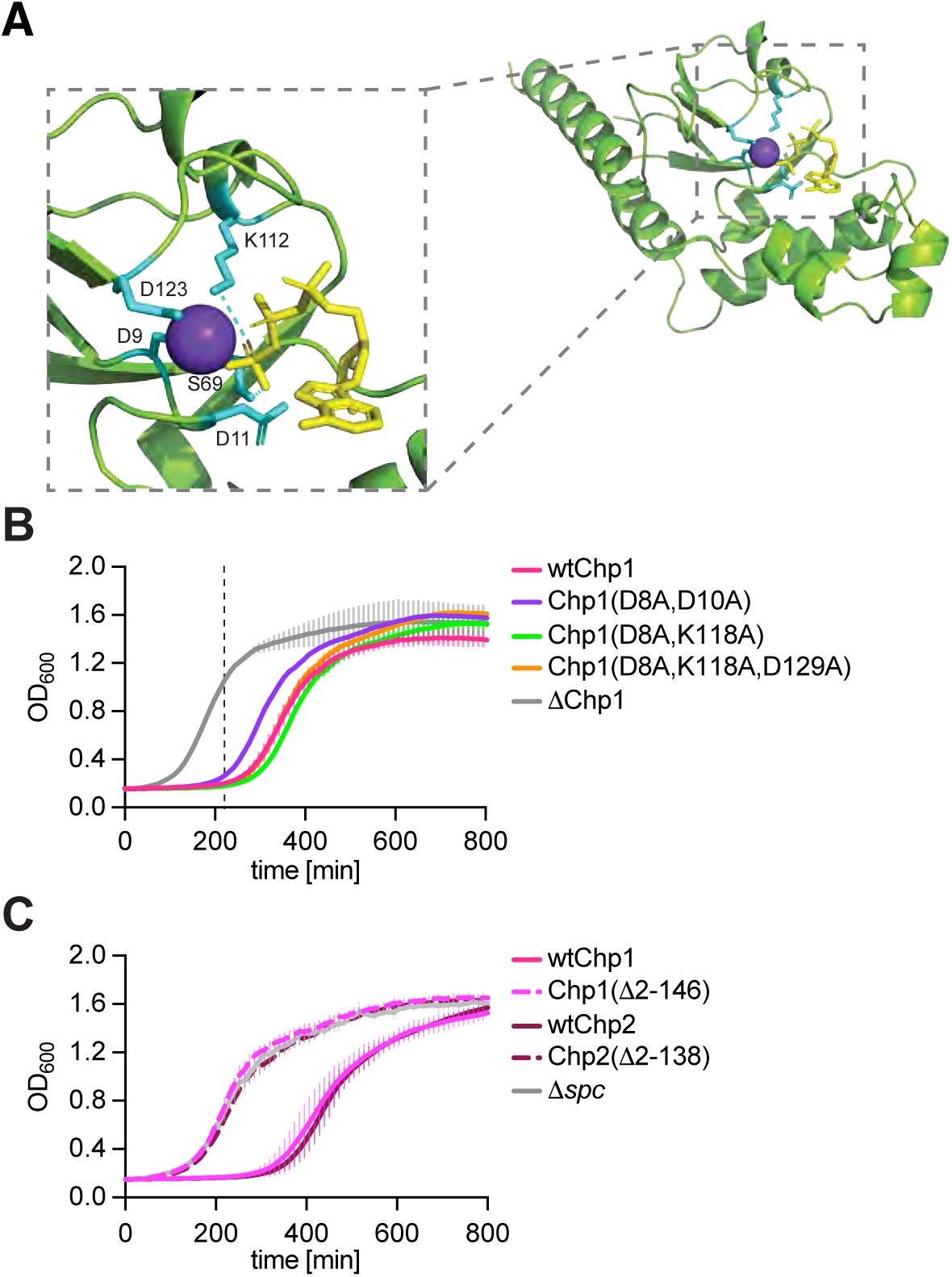
**(A)** Alphafold3 structure of HAD phosphatase domain from Chp2 with conserved residues noted (cyan). Modeled with ATP (yellow) and Mn^2+^ (purple). **(B)** Growth of staphylococci carrying pTarget and pCRISPR dHD with Chp1 variants, measured as OD_600_ after the addition of aTc. Dotted line represents time 220 minutes. Data are mean of two biological triplicates ± SEM. **(C)** Growth of staphylococci carrying pTarget and pCRISPR variants, including deletion of HAD domains in Chp1 and Chp2, measured as OD_600_ after the addition of aTc. Dotted line represents time 220 minutes. Data are mean of two biological triplicates ± SEM.

